# Native epitranscriptome sequencing reveals Hepatitis B virus RNA stability protected by heavy m^6^A modification

**DOI:** 10.1101/2025.04.08.647702

**Authors:** Pei-Yi (Alma) Su, Yun-Hua Lin, Shania Foustine, Chih-Hsu Chang, Ankit Gupta, Brian N Papas, Wiep van der Toorn, Max von Kleist, Redmond P Smyth, Marcos Morgan, Pei-Jer Chen, Kevin Tsai

**Affiliations:** Institute of Biomedical Sciences (IBMS), Academia Sinica, Taipei, Taiwan; Institute of Microbiology, College of Medicine, National Taiwan University, Taipei, Taiwan; Reproductive and Developmental Biology Laboratory, National Institute of Environmental Health Sciences, National Institutes of Health, Durham, NC, USA; Integrative Bioinformatics, Biostatistics and Computational Biology Branch, National Institute of Environmental Health Sciences, National Institutes of Health, Durham, NC, USA; Systems Medicine of Infectious Disease (P5), Robert Koch Institute, Berlin, Germany; Department of Mathematics and Computer Science, Freie Universität Berlin, Germany; Institute of Molecular and Cellular Biology, Strasbourg, France; Hepatitis Research Center, National Taiwan University Hospital College of Medicine, Taipei, Taiwan

**Author notes:** **Authors contributions:** PYS performed most of the experiments, data analysis and wrote the initial draft. YHL performed the STM2457/UZH2-treated HBV-transfection experiments. SF assisted in the RNA half-life assays. CHC assisted with the setup of RNA half-life assays and HBV infection systems. AG, BNP & MM assisted with the DeePlexiCon multiplexing system. WVT, MK & RS assisted with the WarpDemuX multiplexing system. MM initiated the idea of simultaneous detection of m^6^A and polyA tails with Nanopore. KT performed most bioinformatic analysis. PJC & KT supervised the study. RS, MM, PJC & KT obtained funding. KT edited the manuscript. PYS, YHL, AG, BNP, WVT, RS, MM & PJC provided input on the manuscript. **Data availability:** All sequencing data have been deposited at the NIH GEO database under accession number GSE293076.

**Keywords:** RNA, Hepatitis B virus (HBV), N6-methyladenosine (m6A), Nanopore, Virus Replication, RNA Stability, Poly A

## Abstract

Although the RNA of Hepatitis B virus (HBV) is known to be regulated by the epitranscriptomic modification N^6^-methyladenosine (m^6^A), it remains controversial whether if m^6^A is beneficial or detrimental for viral replication. Here, we employ Nanopore direct RNA sequencing (DRS) to analyze the HBV m^6^A epitranscriptome on a single molecule level and elucidate how m^6^A regulates the HBV replication cycle. We found more than 60% of all HBV RNA in cells are m^6^A methylated, ∼3x more methylated than host mRNAs. Previously unreported m^6^A sites were found in *pol* gene, which when mutated reduced viral production. A sizable proportion of HBV RNAs carry poly(A) tails at least twice as long (>120nt) as cellular mRNAs, with the shorter-tailed population more pronounced among m^6^A-free transcripts. In contrast, encapsidated RNAs have shorter poly(A) tails and less m^6^A. Concordantly, pharmacological reduction of viral m^6^A using the m^6^A methyltransferase inhibitor STM2457, shortens viral poly(A) tails and reduces viral RNA stability. Our findings thus suggest that the abundance of m^6^A on viral RNAs protects the surprisingly long viral poly(A) tails and thus enhances RNA stability. Overall, m^6^A strongly enhances HBV viral replication, in support of RNA modification machinery as a promising class of antiviral targets.

## Introduction

Hepatitis B virus (HBV) is a major cause of chronic hepatitis and hepatocellular carcinoma(1). The DNA genome of HBV is reverse transcribed from a pre-genomic RNA (pgRNA)(2), making RNA essential not just for instructing viral protein translation but also for genome replication. RNA is subject to co- and post-transcriptional processes, among which, alternative splicing expands the viral genome coding, with the viral spliced transcript SP1 encoding a core protein variant that contributes to viral entry(3), while RNA polyadenylation promotes translation and acts as a degradation time clock, as transcripts with poly(A) tails <30 nucleotides (nts) are rapidly degraded(4). In addition, small chemical modifications termed epitranscriptomic RNA modifications, have emerged as an important layer of RNA regulation governing splicing, translation, and decay(5).

Multiple RNA modifications are found on viral RNAs at stoichiometries several fold higher than that on the average cellular mRNA, suggesting that viruses have evolved to enrich for modifications that benefit the virus (6–8). Indeed, RNA modifications can support viral gene expression, while in some cases preventing innate immune detection of viral RNA (8,9). Evidence shows that HBV replication is regulated by RNA modifications, with colleagues and us recently demonstrating the cytidine methylation, 5-methylcytidine (m^5^C), essential for viral replication and genomic DNA production (10,11). HBV RNA can also be methylated on the N^6^-position of adenosines (N^6^-methyladenosine, m^6^A). One group showed that m^6^A marks on the 5’ packaging signal (epsilon) promotes viral genome reverse transcription, while m^6^A on the 3’ end of viral transcripts or HBx coding region destabilizes viral RNA and suppresses protein production(12–14). However, a different group instead found m^6^A to support viral gene expression(15). Thus, whether m^6^A enhances or suppresses HBV replication remains controversial.

Early viral epitranscriptomic studies mostly depend on methyl-RNA immunoprecipitation (MeRIP/m^6^A-seq), where m^6^A antibodies were used to pull down m^6^A-containing RNA fragments of ∼100nts, which were subsequently converted to cDNA and sequenced(16,17). While this highly mature strategy has contributed to numerous discoveries, it suffers from low resolution (the exact methylated nucleotide on the 100nt fragment unresolved), high background, and the modification stoichiometry confounded by the reverse transcription (RT) and PCR steps preceding Illumina sequencing(18). Recent advances of Oxford Nanopore direct RNA sequencing (DRS) enable the native long-read sequencing of RNA, detecting canonical as well as modified bases. This allows simultaneous readouts of splice-form-differentiating reads, read abundances free of RT/PCR-biases, RNA modifications, and poly(A) tail lengths, all at the single transcript level(19–21).

Here, we utilized Nanopore DRS with the latest RNA-specific pore to build a comprehensive overview of the HBV m^6^A epitranscriptome. Across HBV replicon-transfected and infected hepatocytes, we found HBV transcripts 3x more likely to be m^6^A-methylated than cellular mRNAs. 13 m^6^A clusters were identified across the viral genome, with a previously unreported cluster on the *pol*-encoding transcript important for viral replication. Curiously, a large proportion of viral transcripts carry poly(A) tails 2x longer than the average ∼60nt tails of cellular mRNA. In contrast, encapsidated viral transcripts are ∼30% less m^6^A-methylated than intracellular viral RNA, with relatively shorter tails. Separating viral transcripts by the presence or absence of m^6^A revealed that m^6^A-free viral transcripts are more likely to possess shorter tails. Concordantly, pharmaceutical reduction of m^6^A levels increased the population of short-tailed viral transcripts, leading to decreased viral RNA stability. We thus argue that overall, m^6^A on HBV RNA maintains their long poly(A) tails, in turn preventing their premature degradation. Lastly, we demonstrate that m^6^A methyltransferase inhibitors confer potent antiviral activity against HBV, in support of RNA modification machinery as a promising class of antiviral targets.

## Materials and Methods

### Cell lines and viruses

HuH-7 and HepG2-NTCP-C4 hepatocytes were generous gifts from Chiaho Shih (Academia Sinica) and Koichi Watashi (National Institute of Infectious Diseases, Japan)(22). The HBV ayw replicon pCHT-9/3091, was used for all experiments unless otherwise specified(23). The Y63D HBV 1.1mer replicon harbors a mutant polymerase with defective reverse transcription priming activity (24). All m^6^A mutant replicons were generated from pCHT-9/3091 by QuickChange Lightning Site-Directed Mutagenesis (Agilent).

### PCR primers and oligonucleotides

All PCR primers and oligonucleotides are listed in Table S3.

### Extraction of HBV viral RNA

Viral particles were concentrated over a 20% sucrose cushion and the RNA extracted with TRIzol, as previously described(11).

### Nanopore direct RNA sequencing (DRS)

Sequencing libraries were prepared using kit SQK-RNA004 (Oxford Nanopore Technologies) using 1 μg of intracellular virion RNA, 50 ng of extracellular virion RNA, or 2 μg of total cellular RNA. DeePlexiCon or WarpDemuX-barcoded adaptors (RTA) were used for multiplex sequencing(25,26). Sequencing was done with FLO-MIN004RA flow cells using a MinION Mk1B (Oxford Nanopore Technologies).

### DRS data analysis

Nanopore output files were demultiplexed using DeePlexiCon or WarpDemuX, with models retrained for RNA004 libraries(25,26). Bases were called using Dorado v0.8.1 with the rna004_130bps_sup@v5.1.0 model, and --modified-bases m6A_DRACH. Reads were aligned to the HBV genome with minimap2 -ax splice -uf -k14, unmapped/reverse/non-primary alignments subsequently removed (samtools view -F 2304). m^6^A peaks were summarized using modkit pileup, filtering for mod calls within AAC/GAC motifs, and peaks with excessive modification no-call events (N_nocall_ > 5%) discarded. The methylation information (MM/ML tags) from Dorado-output bam files were copied onto minimap2-aligned bam files by matching readIDs. Per-read methylation info was extracted using modkit extract --read-calls [calls.file] --ref HBVgenome.fa --motif GAC 1 --motif AAC 1, retaining m^6^A calls with >0.8 probability. PolyA tails were estimated using Dorado with --estimate-poly-a, and the tail length of each read joined to the m^6^A calls table by matching ReadIDs. IsoQuant was used to identify lists of ReadIDs matching each splice form(27), which was then used to bin splice form-matching reads for polyA tail and modification analysis. For analysis of m^6^A1907, HBV-aligned bam files were filtered for alignments over 120nts (samtools view -e ‘(qlen-sclen)>120’), subjected to IsoQuant splice analysis, the uniquely unspliced-assigned ReadID list then used to pull entries from the MM/ML-relabeled HBV-aligned bam files, and summarized with modkit pileup.

The HBV reference genome used is GenBank #J02203.1 reordered with 1818-3182nts forming the first half, joined with 1-1937nts at the end. Host reads were aligned to hg38. HBV known splice form reference as published (28).

### Detection of secreted HBV Virions

Extracellular HBV virions were precipitated using 10% polyethylene glycol (PEG), as previously published (29).

### Northern and Southern blot

Done using DIG-labeled full-length HBV DNA probes as previously described(30).

### Native agarose gel and Western blot of capsid particles

As previously reported(30), capsids were resolved using 1% native agarose gel electrophoresis. Western blots were stained using antibodies: lab-generated rabbit anti-HBc (31), rabbit anti-HBs (Novus NB100-62652), and goat anti-rabbit IgG-HRP (Sigma A6154).

### Detection of HBV pgRNA, Capsid-Associated RNA, and DNA

Encapsidated viral RNA and DNA were quantified via RT-qPCR as previously described(32).

### HBV infection assay

HBV virions were collected from the supernatants of HBV-transfected cells. HepG2-NTCP-C4 cells were cultured in DMEM supplemented with 2.5% DMSO for 72 hours to induce differentiation. The differentiated cells were then infected with HBV at 1,000 genome equivalents (GE) per cell in DMEM containing 2% FBS, 4% PEG-8000, and 2.5% DMSO(22,33).

### ELISA for HBsAg and HBeAg

HBsAg and HBeAg ELISA assays (General Biologicals Corporation, SURASE B-96 (TMB) & EASE BN-96 (TMB)) were done following the manufacturer’s instructions.

### RNA stability assay

HuH-7 cells were transfected with HBV and pulsed with 400μM 4SU 48hrs later. RNA stability then assessed using Roadblock-qPCR(34).

### m^6^A MeRIP-qPCR assay

RNA from HBV-transfected HuH-7 cells or supernatant virions was DNase-treated, fragmented with 10mM ZnCl_2_ in 10mM Tris-HCl (pH 7.0) at 94°C for 45secs, then incubated with m^6^A antibody (Abcam #ab151230) and Dynabeads Protein G beads (Invitrogen) in MeRIP binding buffer (10 mM Tris-HCl, 150 mM NaCl, 0.1% NP-40, pH 7.4) at 4°C overnight. The beads were subsequently washed with MeRIP wash buffer (10 mM Tris-HCl pH 7.4, 1 M NaCl, 0.1% NP-40), and the bound RNA purified using TRIzol. 10% input and m^6^A-enriched RNA were quantified via RT-qPCR.

### Viral replication assays with m^6^A methyltransferase inhibitors

For HBV transfection, HuH-7 cells (1.5 ×10^6^ cells/60-mm dish) were transfected with 5 μg pAAV-HBV1.2-A (genotype A) and 0.5 μg pEGFP-N1 with Lipofectamine 2000 and harvested 7 days later. HBV-transfected or -infected cells were treated with the m^6^A methyltransferase inhibitors STM2457 (Sigma SML3360 or MedChemExpress HY-134836) or UZH2 (MedChemExpress HY-115717), with the inhibitor-containing media refreshed every day for transfected cells or every three days for infected cells. For extracellular protein analysis, supernatants from HBV-transfected cells were concentrated over a 20% sucrose cushion.

### Quantification and statistical analysis

All error bars denote standard deviation (SD). Statistical significances determined by Student’s T-test, *=p < 0.05, **=p < 0.01, ***=p < 0.001.

## Results

### HBV transcripts are highly m^6^A methylated with several previously unreported methylation sites

Our prior Ultra-Performance Liquid Chromatography-Tandem Mass Spectrometry (UPLC-MS/MS) profile of the HBV RNA modification landscape revealed that HBV RNA is enriched with multiple modifications at stoichiometries several-fold higher than cellular mRNAs(11). Among which, m^6^A was found on 4.019% of all HBV adenosines, eight-times higher than host mRNAs, and by far the most abundant of all profiled modifications on HBV RNA (Fig. 1A). Concordantly, m^6^A RNA immunoprecipitation (MeRIP) also showed a high affinity of viral RNA to m^6^A antibodies (Fig. 1B).

**Figure 1.**
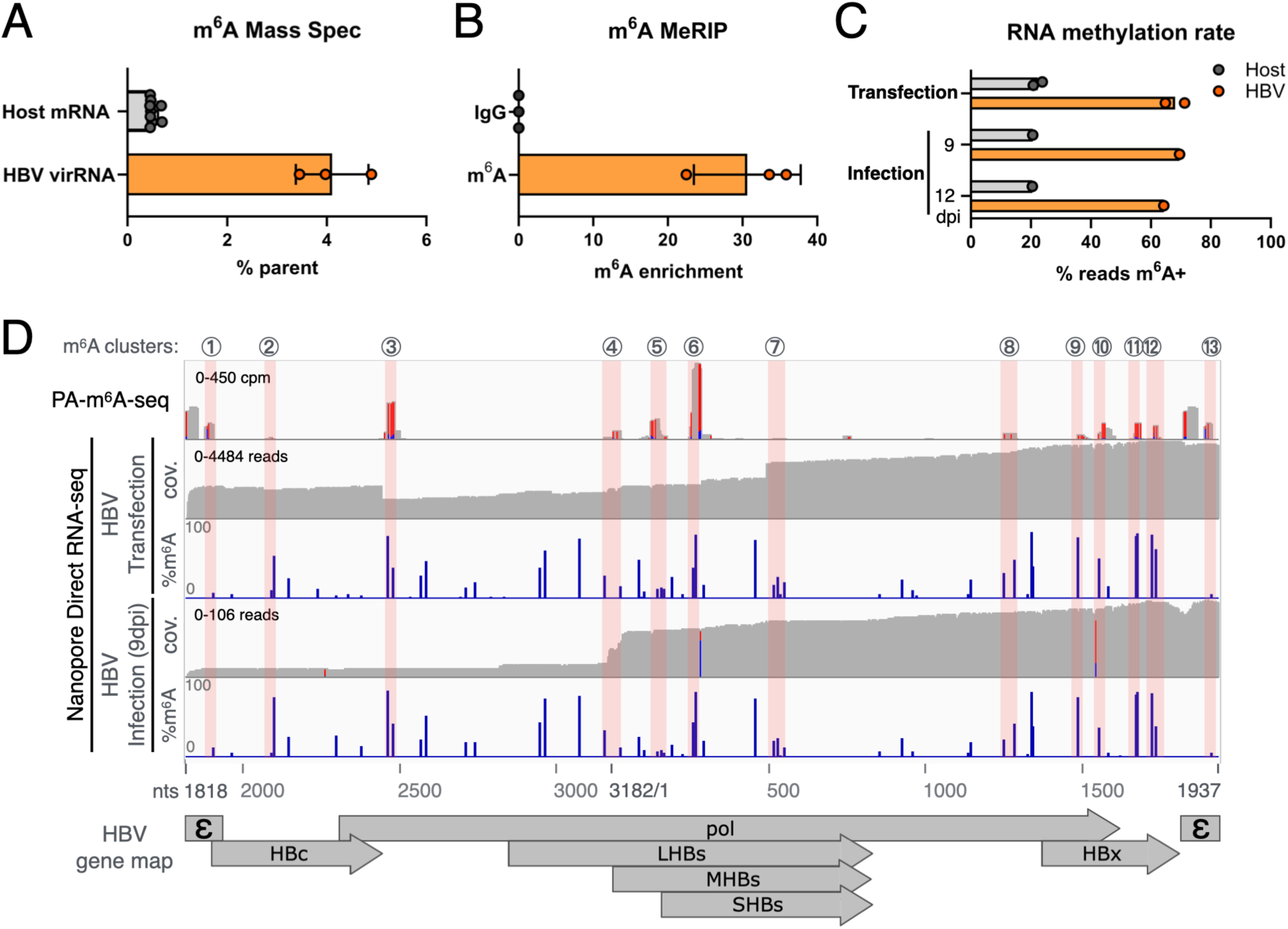
HBV RNA is highly m^6^A methylated. **(A)** Percent m^6^A/A of host and HBV virion RNA quantified by UPLC-MS/MS (adapted from Su et al.(11)). **(B)** HBV RNA m^6^A+ rate assayed by methyl-RNA immunoprecipitation (MeRIP) using a Y63D RT-mutant replicon. **(C)** m^6^A+ rate of host and HBV RNA in replicon-transfected or infected cells, assayed via Nanopore DRS. **(D)** HBV RNA m^6^A sites detected by Nanopore (tracks 2-5), compared to photo-crosslinking-assisted m^6^A sequencing (PA-m^6^A-seq, track 1, adapted Su et al.(11)). Tracks 2&4 show the DRS read coverage (cov.); tracks 3&5 depict the percent coverage m^6^A+. Pink shades highlight m^6^A clusters consistent across PA-m^6^A-seq and DRS runs. Nucleotide coordinates marked here and after are that of Genbank J02203.1.

The recent release of an RNA-specific flow cell has greatly improved the read yield of Nanopore direct RNA sequencing (DRS), while gaining native support for the quantitative detection of m^6^A on each read with single nucleotide resolution. We therefore sought to revisit where m^6^A is located on the HBV genome, and how it impacts its lifecycle. RNA from HBV-transfected HuH-7 cells at 5 days post-transfection, as well as infected HepG2-NTCP-C4 cells at 9 and 12 days post infection (dpi), were subjected to Nanopore DRS (Fig. 1C). Consistent with m^6^A hypermethylation on HBV RNA (Figs. 1A-B), ∼60-70% of viral reads were m^6^A+, while 20% of host reads were m^6^A+. We next analyzed the m^6^A locations of viral reads, and cross-referenced this with previous crosslinking and immunoprecipitation-based photo-activated m^6^A-seq (PA-m^6^A-seq) results(11), revealing 13 clusters of m^6^A peaks consistent across PA-m^6^A-seq and Nanopore sequencing of HBV-transfected and infection samples (Fig. 1D, S1A). Notably, m^6^A clusters 1 and 13 correspond to previously reported m^6^A sites on the 5’ copy of the epsilon element (8-15% DRS reads m^6^A+) and the 3’ epsilon (∼7% reads m^6^A+) (Fig. S1B-C), while several known m^6^A sites on the HBx gene were also detected (Fig. S1C)(12,14,15). Additionally, clusters 2-10 contains multiple previously undetected m^6^A sites (Fig. 1D and S1A).

### Mutational removal of newly detected m^6^A sites reduces HBV particle secretion

To shed light on these newly uncovered m^6^A sites, we introduced point mutations to remove the m^6^A sites with strong Nanopore/PA-m^6^A-seq signals. Specifically, we targeted mutations in the terminal protein (TP) and reverse transcriptase (RT) domain-encoding regions of the *pol* gene (clusters 3 & 6, Figs. 1D & 2A). In the TP domain (cluster 3), the major m^6^A site (A2465) could not be removed without affecting the encoded amino acid, we thus introduced an A2465T mutation that changed the Threonine to the structurally similar Serine (Fig 2B). Two such mutant clones both led to a modest reduction in naked capsid secretion but did not affect HBsAg secretion or HBc protein expression (Fig. 2C-D). The other major m^6^A site (cluster 6) overlaps with the *pol* gene RT domain as well as the HBs genes (Fig. 2E). To separately test m^6^A on the two (*pol* & HBs) genes, we first introduced m^6^A-mutations on a *pol*-start-codon-dead (Δ*pol*) replicon trans-complimented with a wildtype *pol*-expression plasmid(35). With the mutation limited to HBs, we observed a slight reduction in viral production (Fig. S2). Intriguingly, when we left the Δ*pol* replicon intact and instead mutated the *pol*-expression plasmid with m^6^A sites A254 & A265 mutated without amino acid changes (Figs. 2E-F), removal of *pol* m^6^A254 and m^6^A265 significantly reduced infectious particle secretion only when both were removed together, suggesting that near-by m^6^A sites may compensate for each other (Fig. 2G).

**Figure 2.**
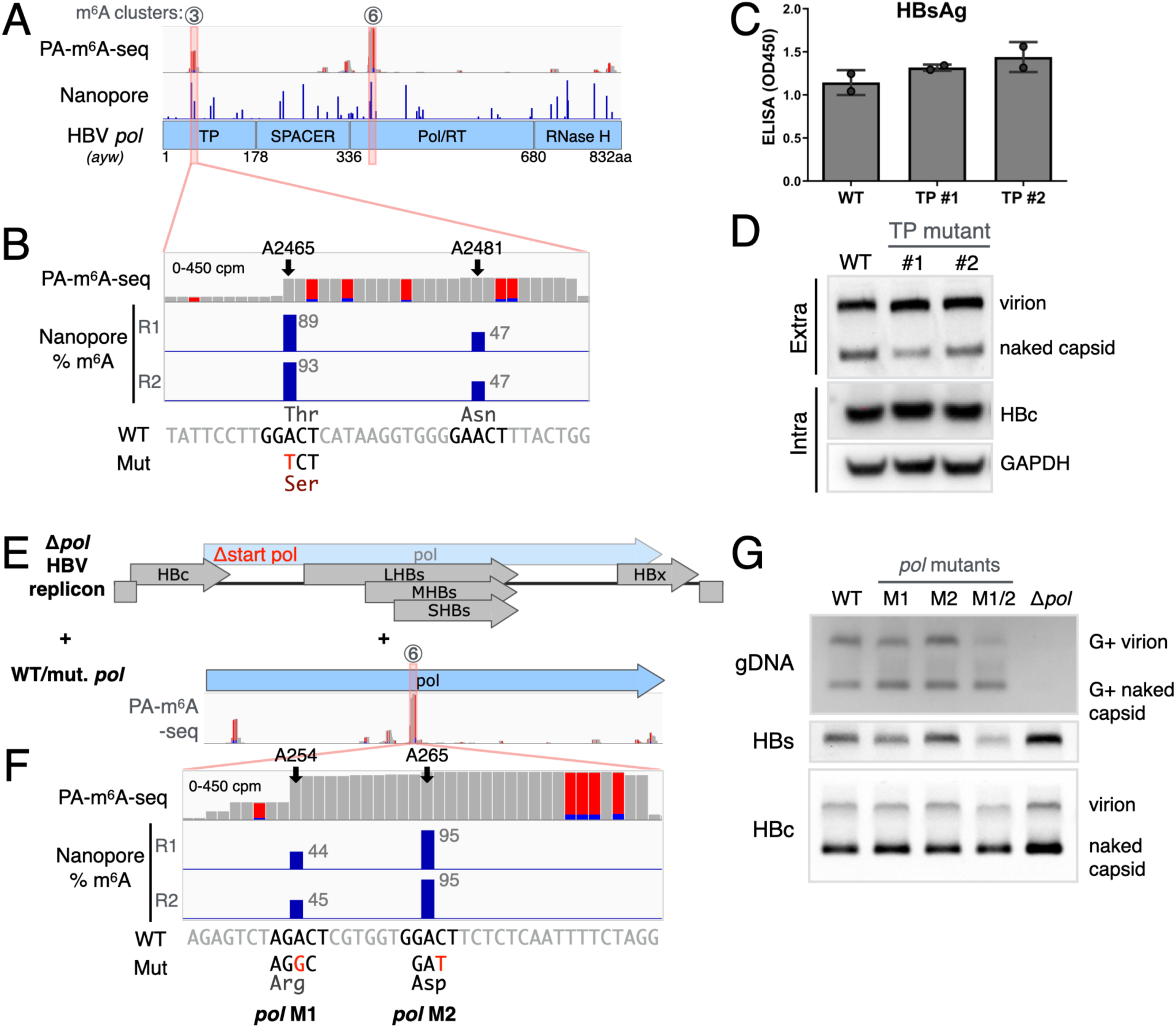
Removal of m^6^A from the *pol* ORF diminished HBV replication. **(A)** m^6^A map on HBV polymerase (*pol*), with m^6^A cluster 3 on TP (Terminal Protein) and cluster 6 on Pol/RT (Polymerase/Reverse Transcriptase) domain. **(B)** Zoomed-in view of m^6^A cluster 3. Bottom tracks depict the wild type (WT) sequence compared with the m^6^A-mutant (Mut) and the encoded amino acid shown bellow. **(C-D)** ELISA measurements of secreted HBs from WT and mutant replicons (A2465T, referred to as TP mutants, clones #1 & #2) **(C)**, secreted (extra) proteins measured via native agarose gel electrophoresis and intracellular (intra) HBc via SDS-PAGE, both followed by Western blotting **(D)**. **(E)** Schematic of Δ*pol* HBV replicon (polymerase start codon deactivated) complemented with WT or m^6^A-mutant HBV *pol*. **(F)** m^6^A cluster 6, zoomed in. Bottom tracks depict the local sequence, with m^6^A-removing mutations (Mut) marked bellow in red. Mutations A254G (M1) and A266T (M2) retains encoded amino acids, marked bellow. **(G)** Secreted virion gDNA and HBs/HBc proteins from WT or *pol* mutants M1, M2 and M1/2 (A254G/A266T dual mutant) assayed by Southern & Western blots at 5 days post-transfection.

### Intracellular HBV RNAs exhibit higher amounts of m^6^A than encapsidated RNA

Since m^6^A was implicated to be necessary for viral RNA encapsidation(13), we next tested whether m^6^A levels differ between intracellular, encapsidated and extracellular viral particle RNA. Capsids were isolated from HBV-transfected HuH-7 cell lysates or supernatants. As the RT-associated RNaseH may degrade encapsidated pgRNA, an additional intracellular capsid set was harvested from a RT mutant (Y63D) replicon. Capsid-extracted RNA was subject to Nanopore DRS and compared with total RNA extracted from HBV-free and HBV-transfected HuH-7 cells. As in Fig. 3A, 20% of host mRNAs were m^6^A+ regardless of presence of HBV, suggesting that HBV doesn’t noticeably change host m^6^A levels. In contrast, total viral RNA in cells displayed a markedly higher 65% reads m^6^A+, compared to 43-48% m^6^A+ in cell encapsidated viral RNAs, and 33% m^6^A+ on supernatant virion RNA (Sup vir). This lower methylation rate on encapsidated RNA was confirmed by MeRIP (Fig. 3B-C).

**Figure 3.**
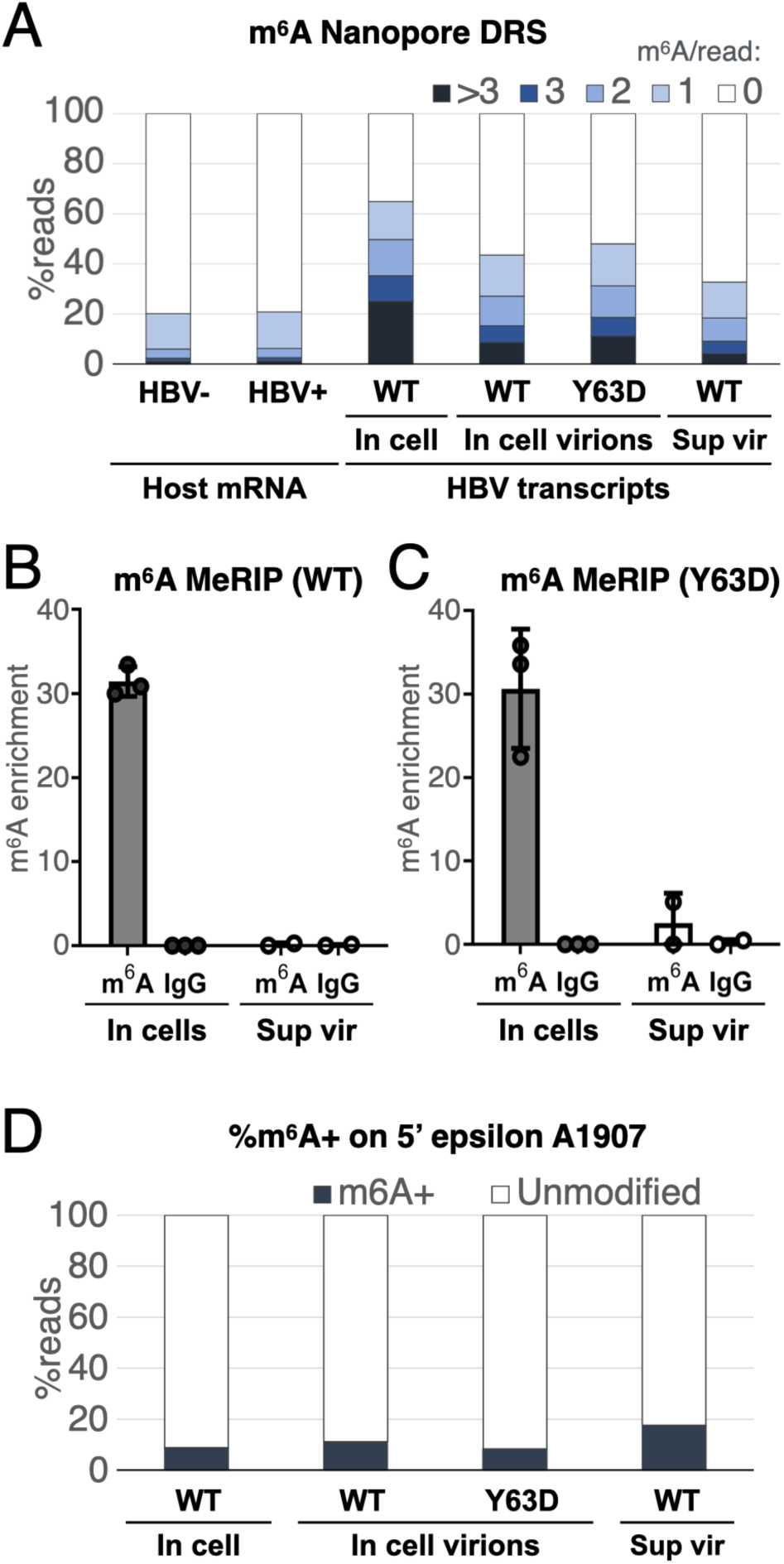
HBV virion-encapsidated RNA are less methylated than intracellular mRNAs. **(A)** Percent of Nanopore DRS reads containing m^6^A, comparing host mRNAs of non-transfected (HBV-) and HBV-transfected (HBV+), as well as HBV-aligning reads of total in cell RNA, in cell core-associated virion RNA (from WT and Y63D RT-mutant HBV), and extracellular virion RNA (Sup vir). **(B-C)** m^6^A methylation rates of intracellular viral RNA and virion-encapsidated RNA in the supernatant (Sup vir) assayed via m^6^A MeRIP on either wild type **(B)** or Y63D RT-mutant HBV **(C). (D)** Percent of reads m^6^A+ among unspliced viral reads covering A1907.

The m^6^A site A1907 on the 5’ epsilon was reported to support encapsidation (12,13). To validate this, we exploited the long read nature of Nanopore to differentiate between methylation sites on the two viral terminal copies of epsilon. To avoid 3’ epsilon reads misaligning to the 5’-copy of epsilon, alignments shorter than one copy of epsilon (120nt) were discarded. Considering that upon encapsidation, it is the unspliced pgRNA that gets reverse transcribed into the genomic DNA, we limited this analysis to unspliced RNA. m^6^A at A1907 was found on 8-11% of reads across in cell and in cell virion datasets, with a slightly higher 17.65% on supernatant virion reads (Fig. 3D). Thus, given the low rate of methylation on A1907, and a minimal difference in %m^6^A between in cell total and encapsidated RNAs, our data do not strongly support a role for m^6^A in pgRNA encapsidation.

### The alternative splicing landscape of HBV

We next analyzed the landscape of HBV alternative splicing. ∼60% of reads could be reliably assigned to a specific isoform, with the remaining reads likely too short to be mapped unambiguously and thus discarded. As in Fig. 4A-D & Table S1, we detected most known HBV splice forms (Fig. S3), with the unspliced pgRNA the predominant RNA species, accounting for 56% of assignable intracellular reads and 65-66% in capsids (Figs. 4A-B, D). Consistent with prior reports, SP1 was the major spliced RNA species in both total HBV RNA and encapsidated viral RNA, representing 20-25% of viral reads in all samples (Figs. 4A-B, D)(3). While the unspliced pgRNA is expected to be the main target of encapsidation, the representation of unspliced RNA in capsids is likely subdued due to RT-associated RNaseH degradation, as the unspliced is as high as 77% in the RT-dead Y63D mutant capsids (Fig. 4C). Spliced viral transcripts are generally less m^6^A+ than unspliced transcripts (Fig. 4E-G), likely due to several m^6^A sites within introns. Yet, less m^6^A was detected in both spliced and unspliced extracellular viral RNA (Fig. 4D), matching the lower overall methylation rate in supernatant virions in Figs. 3A-C. Unexpectedly, we identified an extra short splice form, with almost everything spliced out leaving two epsilons flanking HBx (Bottom lane of Fig. S3, S4A-D & Table S2, this isoform excluded from calculations in Fig. 3). This HBx-isoform (tentatively termed SpX) was present in total intracellular RNA and seems to be efficiently packaged into viral particles, where it constituted nearly 80% of the encapsidated RNA if included in calculations. The presence of SpX was reported, yet not at this high abundance(30,36).

**Figure 4.**
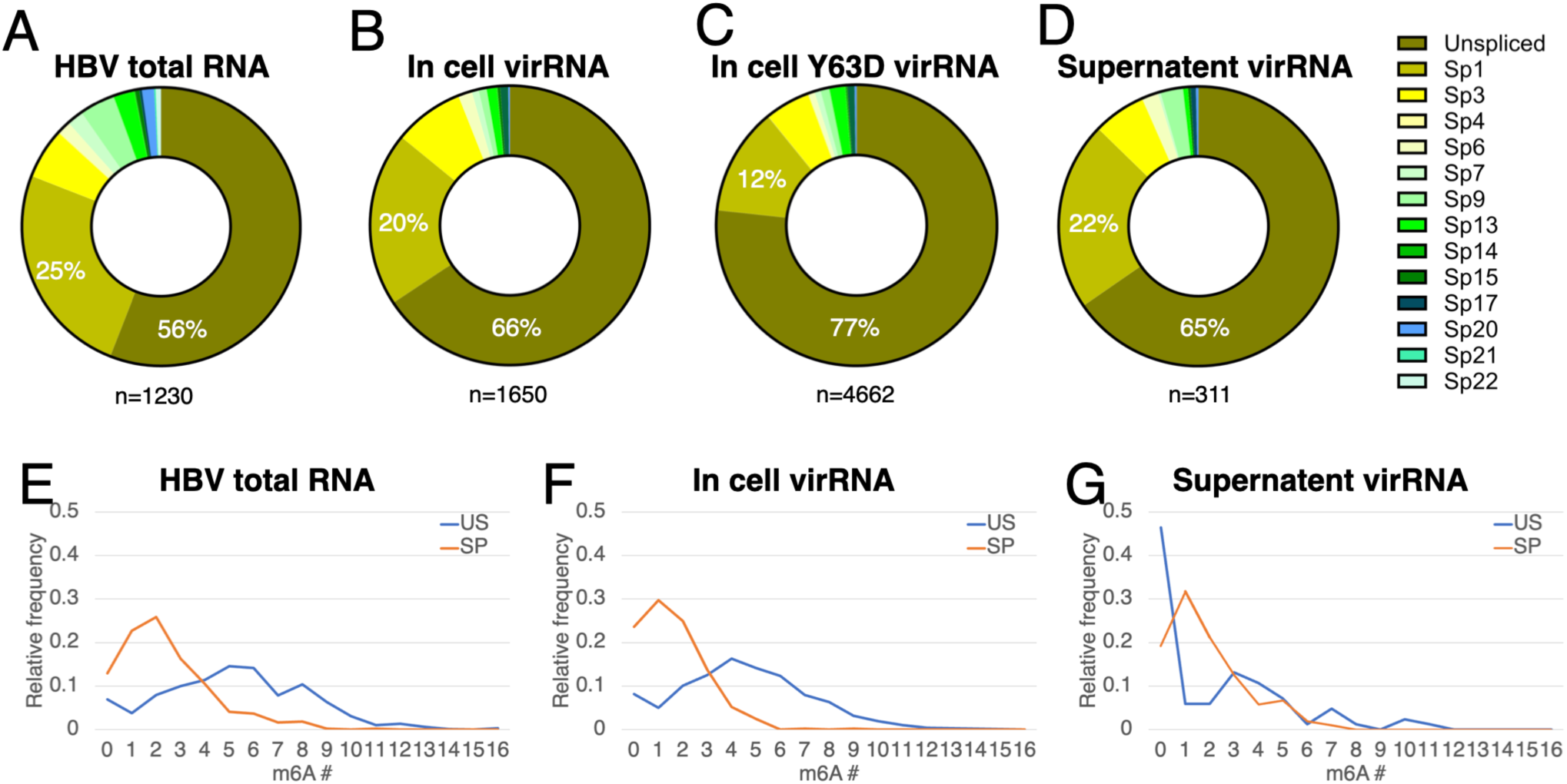
Comparable portions of intracellular and encapsidated HBV RNA are spliced, with spliced RNAs carrying less m^6^A. HBV splice variants were profiled by Nanopore DRS on RNA collected from HBV-transfected HuH-7 cells. The proportions of spliced variants (Fig. S3) are shown for HBV total intracellular RNA **(A)**, intracellular virion-encapsidated RNA **(B-C)**, and extracellular (supernatant) virion RNA **(D)**. **(E-G)** Relative abundances of unspliced (US) or spliced (SP) HBV RNA containing different numbers of m^6^A per transcript, analyzed from total RNA **(E)**, intracellular encapsidated RNA **(F)**, and extracellular RNA **(G)**.

### HBV transcripts harbor extra-long poly(A) tails sustained by m^6^A modifications

RNA poly(A) tails can regulate translation and RNA stability, while m^6^A may regulate poly(A) tails and RNA degradation(4,37), we thus exploited the ability of Nanopore DRS to assay tail lengths and m^6^A status of single transcripts, to elucidate the HBV RNA tails as well as how/if these tails were affected by m^6^A. Upon analyzing human-aligning reads, we do not see a difference in tail length distribution between HuH-7 cells with or without HBV, with a major peak at 60nts (Fig. 5A) In contrast, HBV transcript tails exhibited a bimodal distribution of 60 and 120+nt (Fig. 5B), while encapsidated viral RNA tails were predominantly 60nts (Fig. 5C). Curiously, the poly(A) tails of HBV-infected HepG2-NTCP cells were again ∼60nt for host transcripts, yet viral transcripts possessed exceptionally long poly(A) tails of 150-210nt (Fig. 5D & S5A). When we defined >120nt tails as long tails, we estimate the long tailed proportion of host transcripts at 40%, yet almost 80% of viral transcripts are long tailed (Fig. 5E, S5B). Upon computationally separating reads with or without m^6^A, m^6^A-free viral reads displayed a more pronounced 60nt peak, with an accompanying depletion in the ∼270nt range (Fig. 5F), suggesting that m^6^A is linked to poly(A) tail lengths.

**Figure 5.**
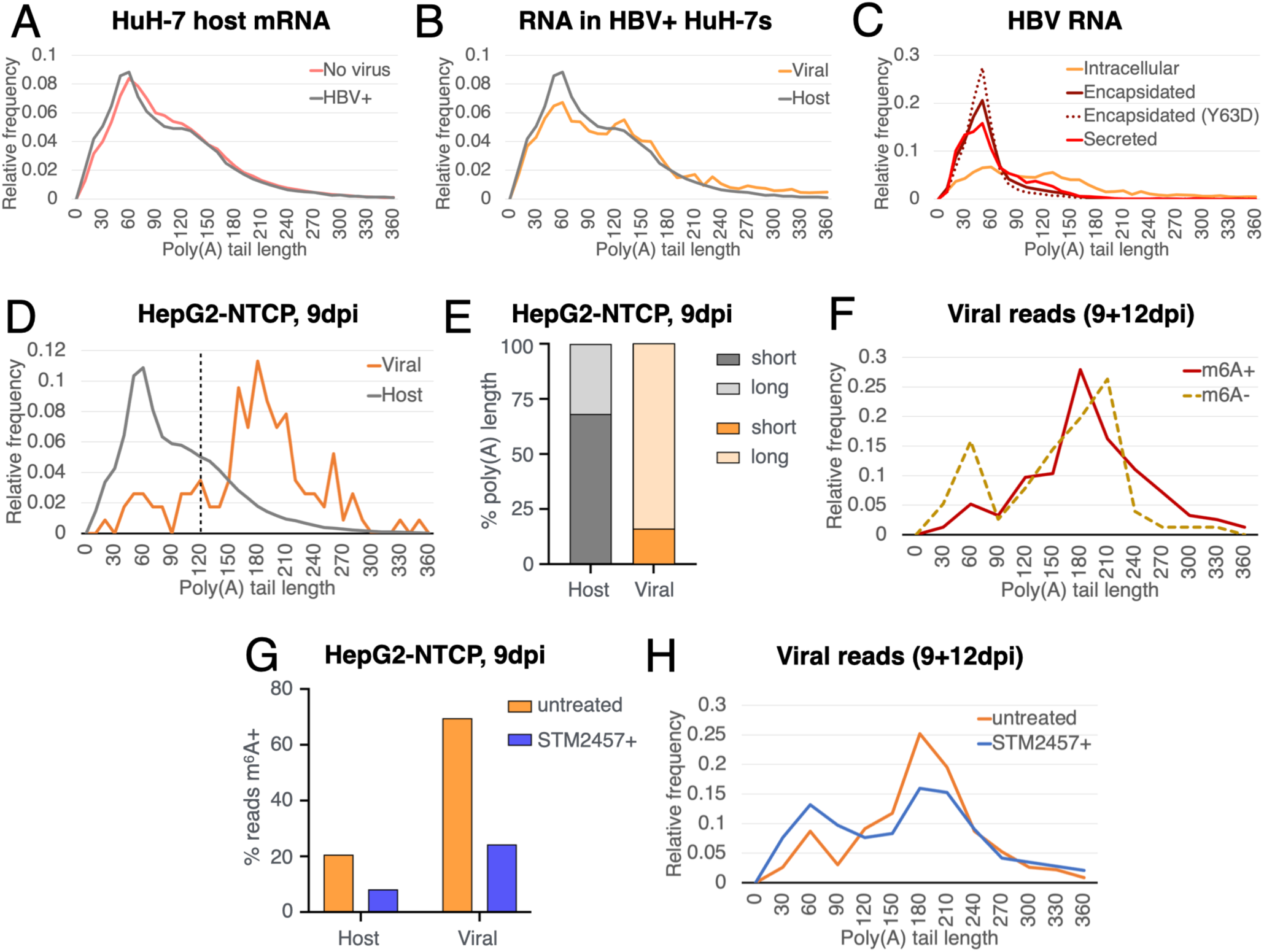
A large proportion of HBV RNAs contain poly(A) tails >60nt, with the presence of m^6^A maintaining longer tails. **(A-C)** Poly(A) length distributions of HBV-transfected HuH-7 RNA, with comparison between untransfected & HBV-transfected host mRNAs **(A**), intracellular host and viral RNAs **(B)**, and intracellular, encapsidated and secreted viral RNA **(C). (D)** Host & viral poly(A) tail lengths in HBV-infected HepG2-NTCP cells, 9 days post infection (dpi). (See Fig. S5 for 12dpi, no tail distribution difference observed between 9 & 12dpi). **(E)** Proportion of transcripts with short (<120nt) and long (>120nt) poly(A) tails in panel D. **(F)** Tail length distributions of m^6^A+ and m^6^A-viral reads **(G)** Percent of m^6^A+ host & viral Nanopore DRS reads with and without treatment with the m^6^A methyltransferase inhibitor STM2457. **(H)** Tail length distributions of untreated versus STM2457-treated HBV-infected cells (9 & 12dpi data combined for increased read counts in panels F & H).

We next tested whether m^6^A regulates poly(A) tail length in viral RNA, hypothesizing that m^6^A removal would lead to the accumulation of shorter RNA tails. HBV infections were therefore repeated in the presence of a well characterized m^6^A inhibitor, STM2457, which binds the primary mRNA m^6^A methyltransferase METTL3 with high specificity(38). STM2457 led to a ∼3-fold reduction in m^6^A+ read calls in both host and viral RNA (Fig. 5G & S5C), coinciding with a diminished 180+nt peak and a more pronounced 60nt shoulder in the poly(A) tail distribution (Fig. 5H). Thus, these results suggest that the high amounts of m^6^A on HBV RNA correlates with long poly(A) tails.

### Pharmacological inhibition of m^6^A methylation suppresses HBV replication by reducing viral RNA levels

As poly(A) tails can protect against RNA degradation pathways that are activated following deadenylation, we next hypothesized that removal of m^6^A may lead to viral RNA instability, and thus an overall decrease of viral gene expression and replication. We thus proceeded to test this using two orthogonal m^6^A methyltransferase inhibitors, STM2457 and UZH2(39). In HBV replicon-transfected cells, both STM2457 and UZH2 lead to a loss of secreted viral proteins, intracellular viral proteins, as well as viral RNA, with STM2457 showing dose-dependent inhibition of viral production (Figs. 6A-C). This suggests that m^6^A enhances viral gene expression on the RNA level, which is upstream of protein production and subsequent encapsidation and reverse transcription. We next tested if m^6^A enhances viral RNA stability, using Roadblock-qPCR where incorporation of 4-thiouridine (4SU) into nascent RNA restricts subsequent RT-qPCRs to only detect RNAs produced prior to 4SU pulse(40). Monitoring the levels of pre-pulse residual RNA overtime revealed that STM2457-inhibition of m^6^A methylation indeed decreased the stability of viral RNAs (Fig. 6D). The antiviral effect of STM2457 was also seen in HBV-infected HepG2-NTCP cells, where STM2457 treatment led to a 50% decrease in viral RNA Nanopore read counts at both 9 and 12 dpi, whereas this reduction was < 10% for host cellular reads. (Fig. 6E). An ELISA analysis for viral antigen secretion from these infected cells confirmed that STM2457 lead to a dose-dependent decrease of HBs and HBe antigens over a 12-day infection course (Figs. 6F-G). We thus conclude that m^6^A hypermethylation on HBV RNA enhances viral replication through maintaining long poly(A) tails on the viral transcripts, which in turn maintains the longevity of viral RNAs.

**Figure 6.**
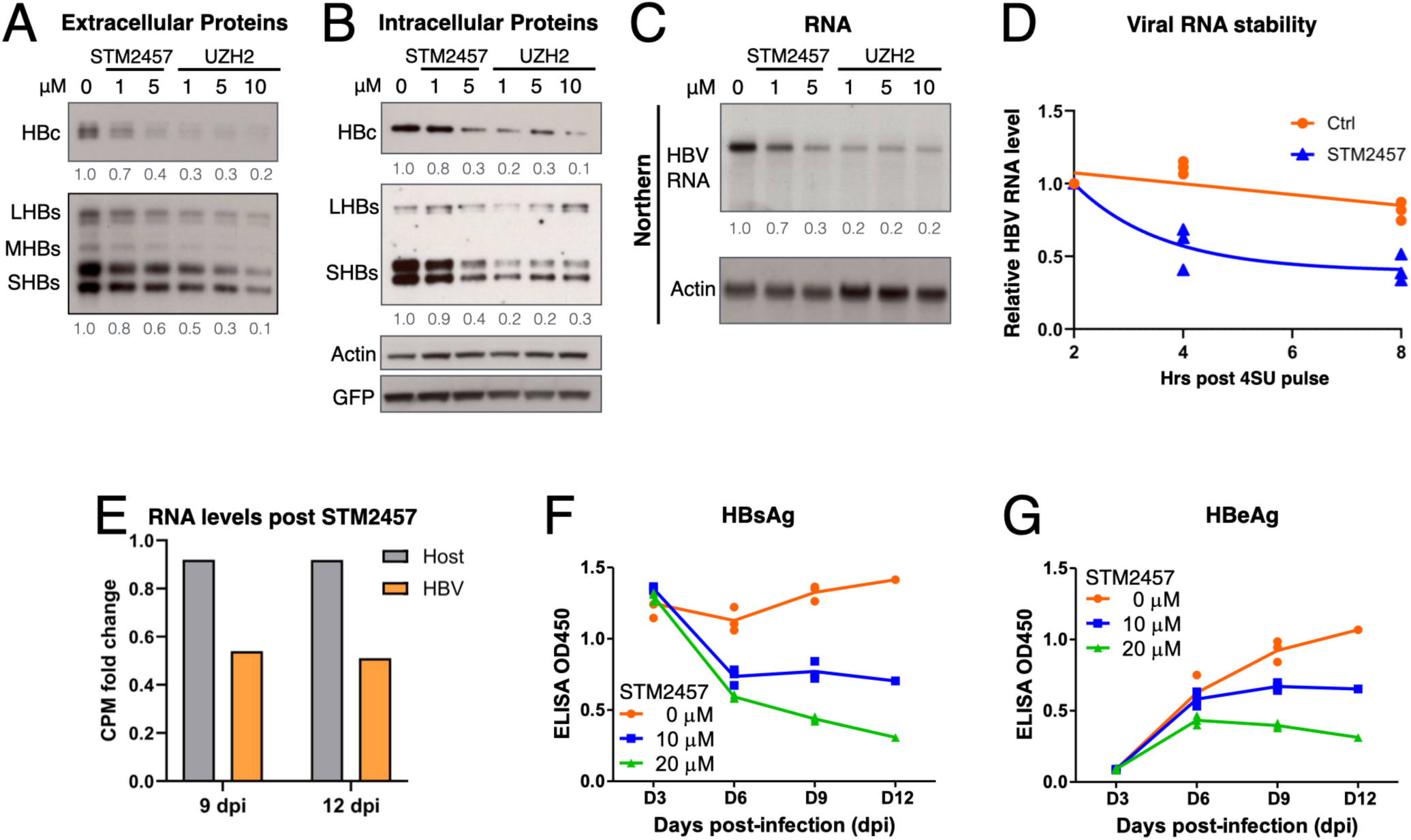
Pharmacological inhibition of m^6^A methylation diminishes HBV replication via reduction of viral RNA stability. **(A-D)** HBV-transfected HuH-7 cells treated with the m^6^A methyltransferase inhibitors STM2457 or UZH2 subjected to Western blot assay of secreted viral proteins **(A)**, intracellular viral proteins **(B)**, Northern blot assays of viral RNA levels **(C)**, and viral RNA stability tested by Roadblock-qPCR **(D)**. **(E)** Nanopore read counts of host or HBV reads from HBV-infected HepG2-NTCP treated with STM2457. **(F-G)** Supernatant HBsAg and HBeAg in HBV-infected cells treated with STM2457 measured by ELISA.

## Discussion

Akin to several other human pathogenic viruses, HBV RNA is methylated with m^6^A, which also impacts viral replication(12–15). This is supported by our recent UPLC-MS/MS quantification of HBV RNA modifications(11), which together with the Nanopore data here, paints a picture of a heavily m^6^A-methylated viral transcriptome that is 8x more methylated by mass and 3x higher in terms of proportion of transcripts m^6^A+, when compared to host cellular mRNAs (Fig. 1).

All prior HBV m^6^A maps have relied on MeRIP/m^6^A-seq. In contrast, we recently performed PA-m^6^A-seq on HBV RNA, which utilizes 4SU-mediated antibody-RNA crosslinking and RNase-footprinting to improve the signal-to-noise ratio of traditional MeRIP/m^6^A-seq, resulting in sharper peaks and the ability to computationally remove background reads lacking crosslink-associated T>C mutations(11,41). The PA-m^6^A-seq data corroborates well with our Nanopore DRS, with 13 consistent methylation clusters identified (Fig. 1D, S1A). Importantly, our PA-m^6^A-seq map informed our need to only accept Nanopore m^6^A calls within the canonical RAC motif, thus greatly reducing the number of questionable modification calls. Several of our identified m^6^A sites match previously characterized sites, including the two epsilon sites, as well as four sites within HBx (12,14,15) (Figs. S1B-C). Here, we found nine m^6^A clusters that seemingly eluded prior MeRIP attempts, among which two m^6^A sites on the *pol* gene was not only within the highest PA-m^6^A-seq peak, but also proved crucial for virion production (Fig. 2). These findings highlight the importance of cross-referencing modification maps from multiple mapping techniques, as each method may have their own inherent biases.

m^6^A on A1907 within the HBV 5’ epsilon element was particularly reported to be required for viral RNA packaging (12,13). We indeed detected m^6^A1907 (Figs. 1D, S1B-C), however this was a rather small peak in the semi-quantitative PA-m^6^A-seq results while detected by Nanopore on 8-9% of reads. If m^6^A1907 were required for encapsidation, encapsidated viral RNA should be much more methylated than the total intracellular pool of viral RNAs. While this was reportedly seen by MeRIP-qPCR(13), we could not repeat this MeRIP result (Figs. 3B-C), and Nanopore detection of m^6^A1907 on encapsidated RNA was only up to 17.65% even when we limited our analysis to unspliced RNA (Fig. 3D). We thus consider that if m^6^A1907 can enhance HBV RNA encapsidation, it would play a minor role. In contrast, when taking all m^6^A sites into account, we found encapsidated and secreted RNA less methylated than intracellular viral transcripts, albeit still higher than host mRNAs (Fig. 3A-C). Whether if this methylation rate difference between intracellular/encapsidated transcripts were a result of packaging selection, RT/RNaseH-associated removal or some other unforeseen factor would require further validation. However, given that the methylation rates between wildtype and the RT-defective Y63D encapsidated RNA were quite close (Fig. 3A), we believe that RT/RNaseH is unlikely to explain this observation.

Of particular interest is the observation that a large proportion of HBV RNA contain poly(A) tails much longer than host mRNAs (Fig. 5B). In replicon-transfected HuH-7 cells, viral RNA tails display a 60/120+nt bimodal distribution as well as a larger distribution of tails in the 240+nt range, which matches that previously reported for HBV RNA in the HBV stable line HepG2.2.15(42). As the >60nt tailed population was not seen in encapsidated or secreted viral RNAs, the extra-long poly(A) tails are likely only needed within host cells (Fig. 5C). This makes sense as the two primary functions of poly(A) tails, defense against RNA degradation and enhancement of translation, both involve cellular machinery that are unlikely to be present in capsids. This also coincides with the observation that pgRNA poly(A) tails are dispensable for encapsidation and the subsequent reverse transcription(43). Curiously, we found HBV RNA tails to be even longer in infected HepG2-NTCP cells, with the majority of tails between 150-210nt (Fig. 5D-E, S5A-B). We consider infection systems a better representation of actual infections than transfection systems, yet both systems show viral RNA carrying extraordinarily long tails and an abundance of m^6^A. Intriguingly, analysis of tail length and m^6^A status on a single transcript/read basis revealed that unmethylated RNA are more likely to have shorter ∼60nt tails (Fig. 5F), which corroborates with STM2457-depletion of m^6^A also increasing the short tail population (Fig. 5G-H). Notably, m^6^A was reported to promote deadenylation and degradation of host mRNAs, yet we previously found m^6^A to instead stabilize HIV-1 RNA. Our finding here that m^6^A protects the poly(A) tails and RNA stability of HBV RNA, suggest a convergence of viral m^6^A function between the two viruses(37,44).

The accumulation of m^6^A on viral RNA is likely evolutionarily beneficial. We thus propose that the high m^6^A likely enhances viral replication by protecting the extended poly(A) tails, thus enhancing RNA stability. Indeed, mutational removal of m^6^A cluster 6 diminishes viral production (Fig. 2E-G), while m^6^A-depletion with STM2457 and UZH2 both lead to diminished viral RNA and virion production, with STM2457 decreasing RNA stability (Fig. 6). One prior study using siRNA-based m^6^A-depletion suggested that m^6^A instead promotes HBV RNA degradation(12), yet a competing siRNA m^6^A-depletion study showed m^6^A enhancing viral RNA levels, in concordance with our results(15). Nonetheless, exploitation of m^6^A to enhance viral RNA expression has emerged as a common theme across DNA, RNA and retroviruses(8,9), with STM2457 antiviral against HBV (Fig. 6) as well as the coronaviruses OC43 and SARS-CoV-2(45). It is clear that HBV goes to lengths to protect their RNA tails(42), and it would be interesting if this m^6^A-poly(A) linkage is a common mechanism utilized by other m^6^A-exploiting viruses as well.

## Acknowledgements

We thank Koichi Watashi for providing us with HepG2-NTCP cells. We also thank the core facilities at Academia Sinica: IBMS DNA Sequencing (AS-CFII-111-211), IMB bioinformatics core and Academia Sinica grid-computing center (ASGC). This research was supported by the NSTC grants 112-2320-B-001-023 to K.T., 110-2314-B-002-034-MY3 and 113-2314-B-002-309-MY3 to P.J.C., Academia Sinica career development award AS-CDA-112-L02 to K.T., the Intramural Research Program of the NIH (project ZIA ES103355), and the Center for Structural Biology of HIV-1 RNA, funded by NIAID, NIH grant U54 AI17660 to K.T. and R.P.S.

---

**Figure S1.**
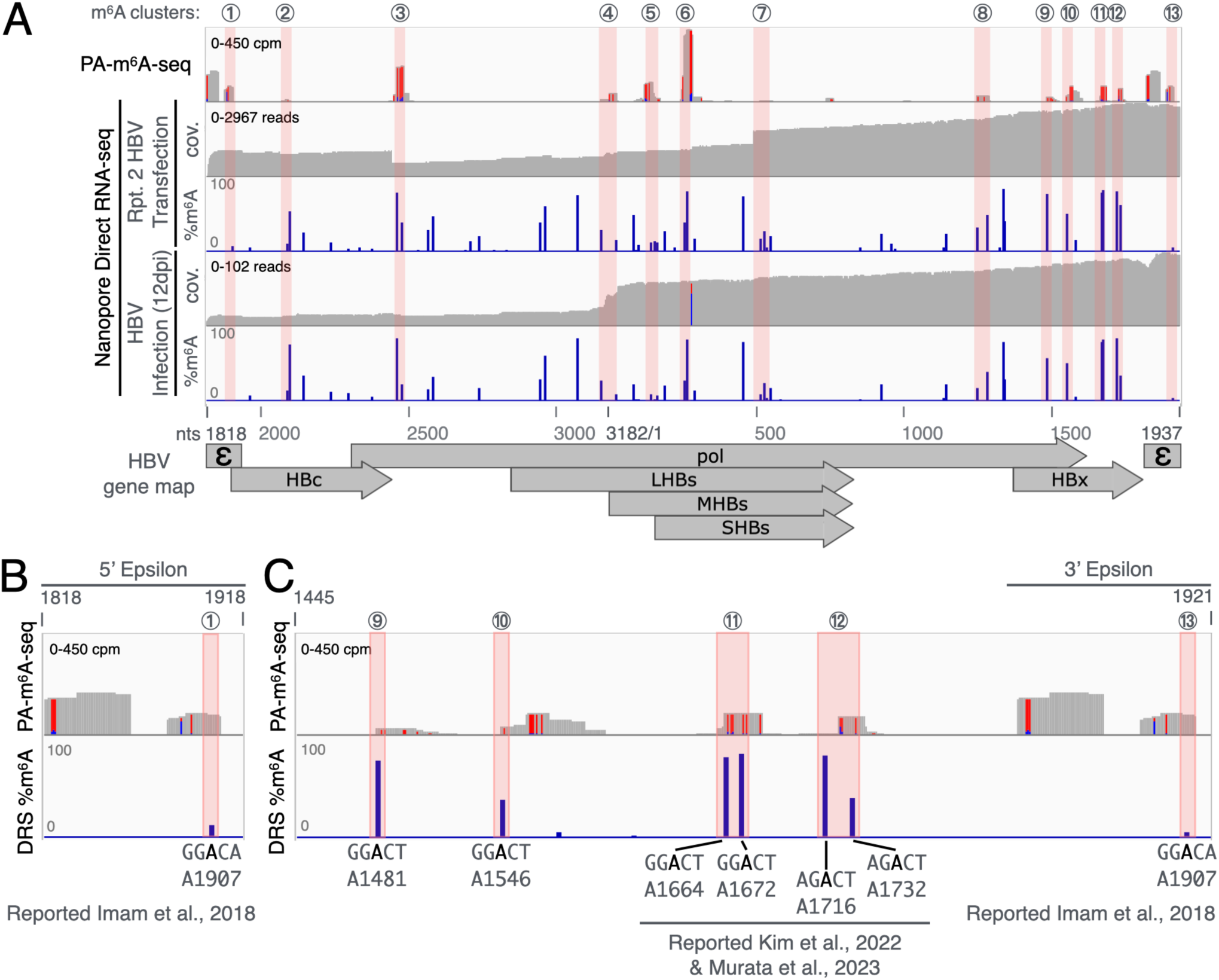
Mapped HBV RNA sites are repeatable with several sites consistent with prior reports. **(A)** HBV RNA m^6^A sites detected by Nanopore DRS (tracks 2-5), compared to previous PA-m^6^A-seq results(11). Tracks 2&4 show the read coverage (cov.) of DRS runs on replicon-transfected HuH-7s (2nd repeat) and HBV-infected HepG2-NTCP cells harvested 12dpi, respectively; tracks 3&5 depict the percent coverage called as m^6^A+ on the DRS runs. Horizontal pink boxes highlight the thirteen m^6^A peaks consistent across PA-m^6^A-seq and all DRS runs (see Fig. 1C for transfection repeat 1 and 9dpi results). **(B-C)** Zoomed view of PA-m^6^A-seq and DRS-detected m^6^A sites (9dpi, Fig. 1C) on the 5’ Epsilon region **(B)**, and 3’ end region up to the 3’ epsilon **(C)**. Nucleotide numbers marked according to Genbank J02203.1 which only has one copy of epsilon, thus both 5’-& 3’-epsilons share the same coordinates.

**Figure S2.**
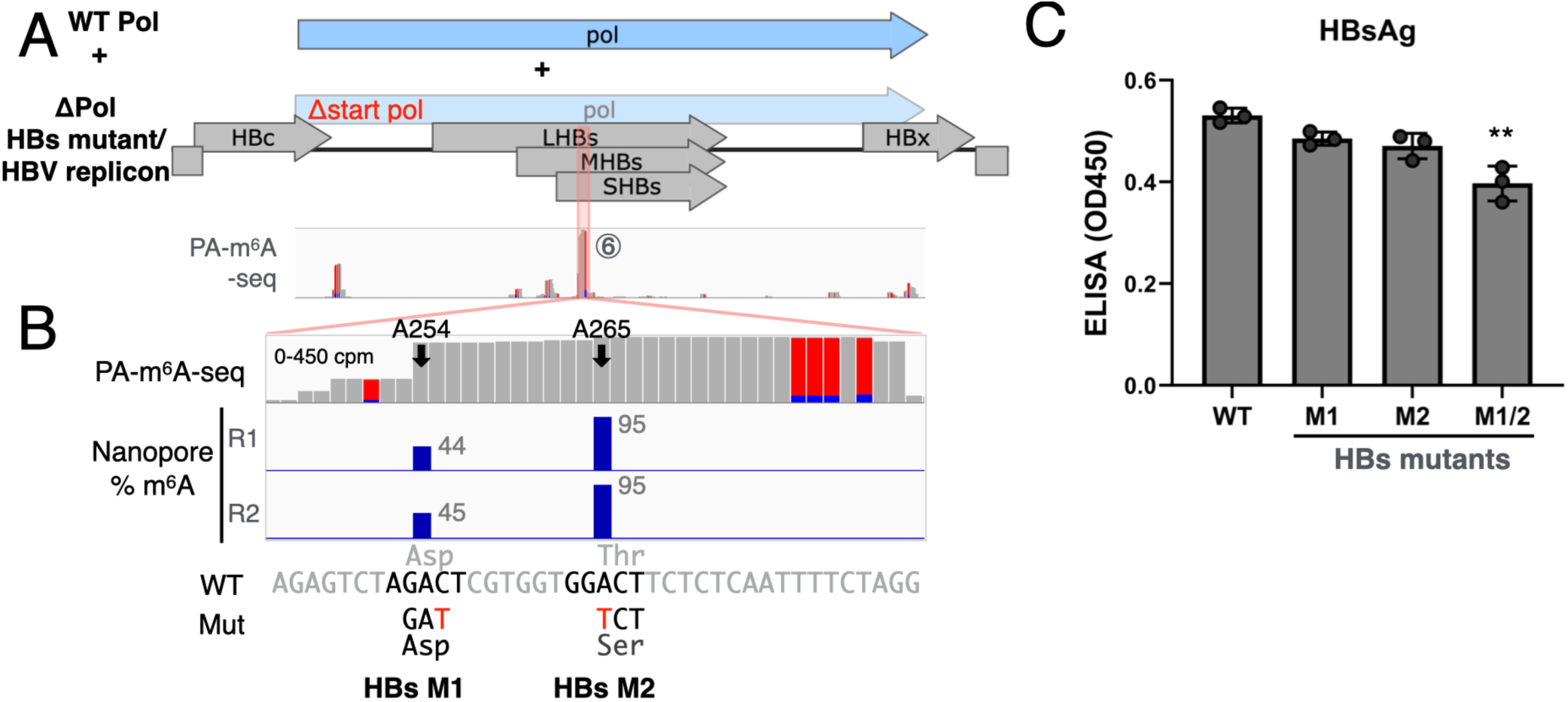
m^6^A254 and m^6^A265 on HBs transcripts mildly affect viral replication. **(A)** Schematic of m^6^A sites on HBs transcripts. **(B)** Zoomed in view of cluster 6, which contains m^6^A254 and m^6^A265. Bottom lanes depict m^6^A-removing mutants with the changed nucleotides in red and the resulting amino acid shown below. **(C)** WT and HBs m^6^A-mutant replicons were transfected into HuH-7 cells, with the secreted virion HBs measured via ELISA. Error bars = SD, n = 3; **P < 0.01.

**Figure S3.**
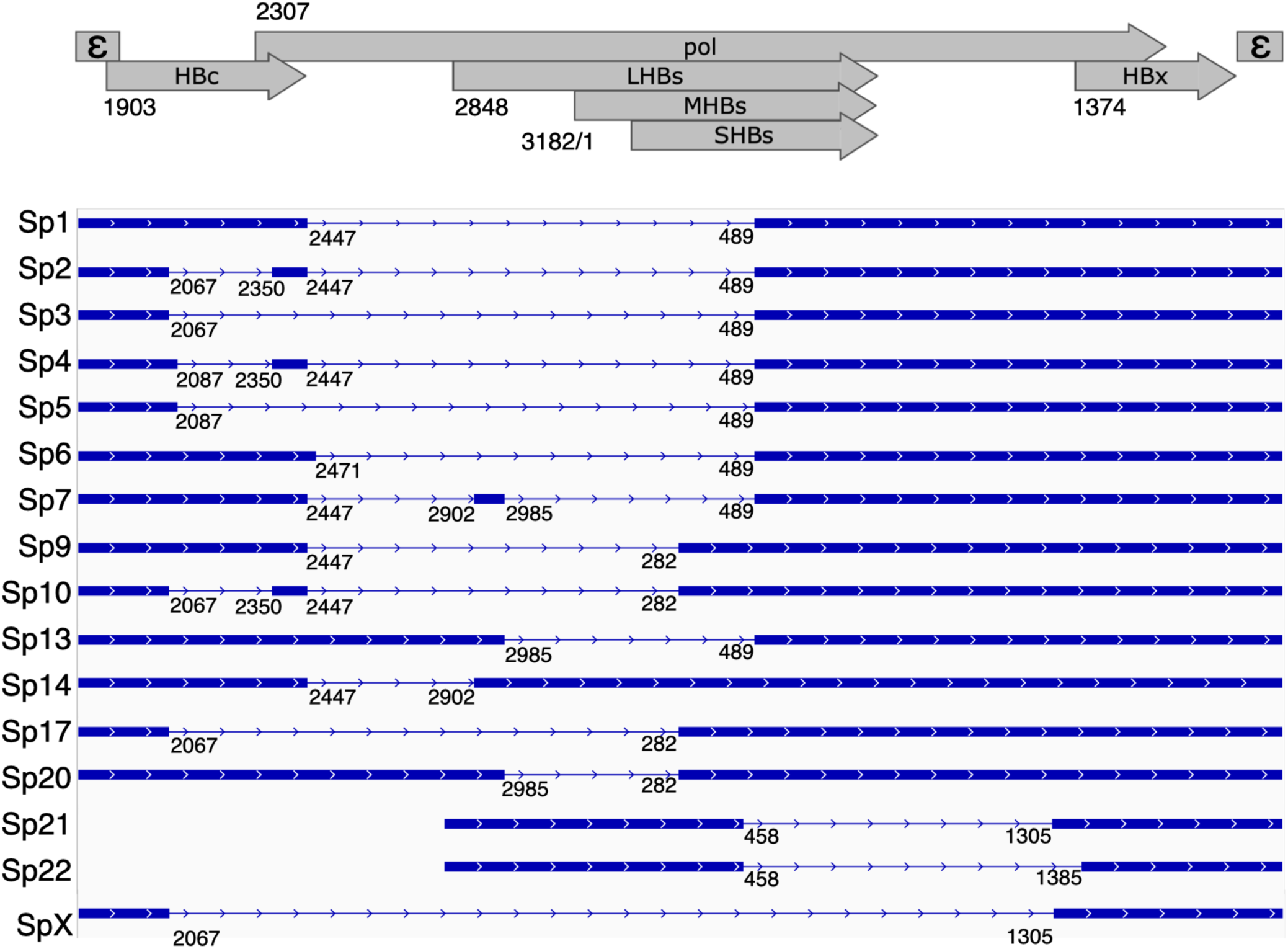
HBV spliced variants detected by Nanopore DRS. Detected HBV spliced variants with their respective splice donor and acceptor sites. Sp1-Sp22 adapted from Kremsdorf et al.(28), with the last isoform (SpX) detected from DRS runs on HBV transfected HuH7 cells.

**Figure S4.**
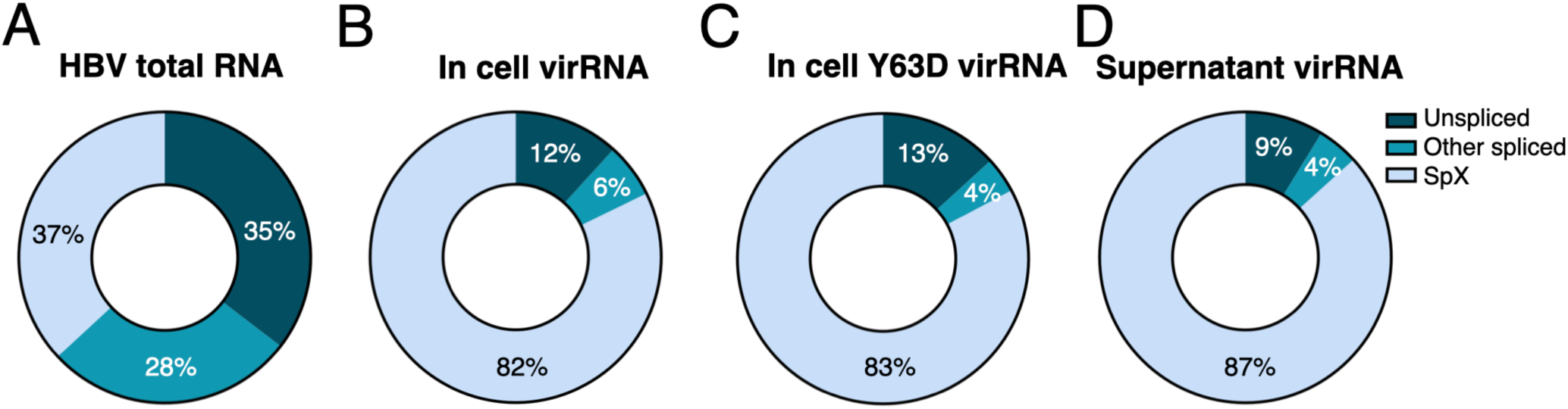
High amounts of the short SpX splice form in HBV viral particles. **(A-D)** The proportion of SpX and other spliced variants derived from HBV-transfected viral RNA shown for intracellular total RNA **(A)**, intracellular encapsidated RNA (WT & Y63D RT mutant) **(B-C)** as well as extracellular (supernatant) viral RNA **(D)**.

**Figure S5.**
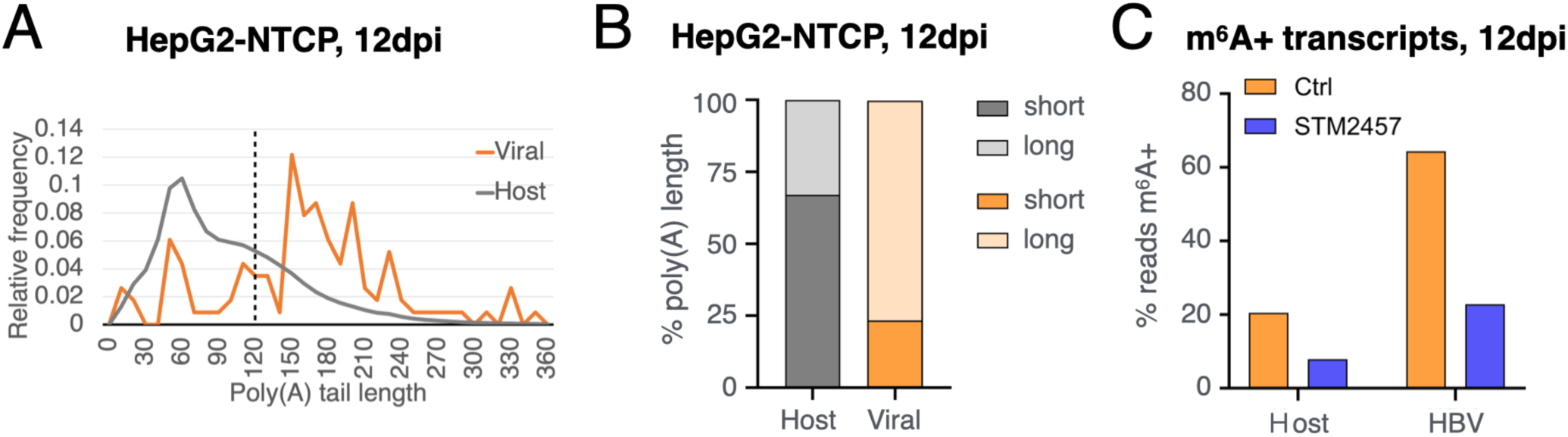
RNA Poly(A) tail length and m^6^A representation in HBV-infected cells, 12dpi. **(A)** Poly(A) tail length distribution of host and viral transcripts in HepG2-NTCP cells 12 days post-HBV infection. **(B)** Proportion of transcripts with short (<120 nt) and long (>120 nt) poly(A) tails in host and viral RNA. **(C)** Percent of DRS reads m^6^A+ host and viral transcripts from infected cells treated with or without STM2457.

**Table S1.**
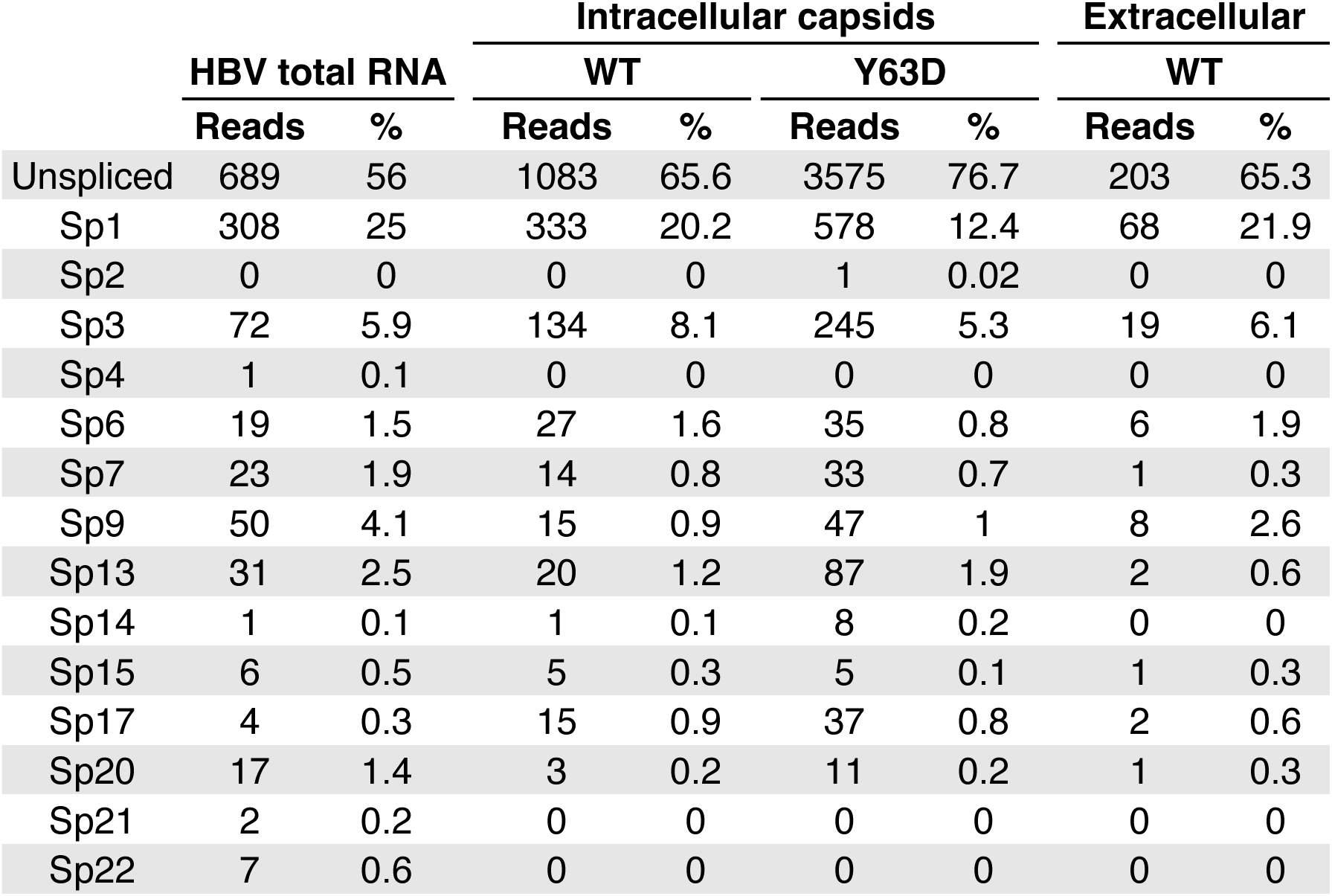
Proportion of HBV spliced variants detected by Nanopore DRS.

|  | HBV total RNA |  | Intracellular capsids |  |  |  | Extracellular |  |
| --- | --- | --- | --- | --- | --- | --- | --- | --- |
|  |  |  | WT |  | Y63D |  | WT |  |
|  | Reads | % | Reads | % | Reads | % | Reads | % |
| Unspliced | 689 | 56 | 1083 | 65.6 | 3575 | 76.7 | 203 | 65.3 |
| Sp1 | 308 | 25 | 333 | 20.2 | 578 | 12.4 | 68 | 21.9 |
| Sp2 | 0 | 0 | 0 | 0 | 1 | 0.02 | 0 | 0 |
| Sp3 | 72 | 5.9 | 134 | 8.1 | 245 | 5.3 | 19 | 6.1 |
| Sp4 | 1 | 0.1 | 0 | 0 | 0 | 0 | 0 | 0 |
| Sp6 | 19 | 1.5 | 27 | 1.6 | 35 | 0.8 | 6 | 1.9 |
| Sp7 | 23 | 1.9 | 14 | 0.8 | 33 | 0.7 | 1 | 0.3 |
| Sp9 | 50 | 4.1 | 15 | 0.9 | 47 | 1 | 8 | 2.6 |
| Sp13 | 31 | 2.5 | 20 | 1.2 | 87 | 1.9 | 2 | 0.6 |
| Sp14 | 1 | 0.1 | 1 | 0.1 | 8 | 0.2 | 0 | 0 |
| Sp15 | 6 | 0.5 | 5 | 0.3 | 5 | 0.1 | 1 | 0.3 |
| Sp17 | 4 | 0.3 | 15 | 0.9 | 37 | 0.8 | 2 | 0.6 |
| Sp20 | 17 | 1.4 | 3 | 0.2 | 11 | 0.2 | 1 | 0.3 |
| Sp21 | 2 | 0.2 | 0 | 0 | 0 | 0 | 0 | 0 |
| Sp22 | 7 | 0.6 | 0 | 0 | 0 | 0 | 0 | 0 |

**Table S2.**
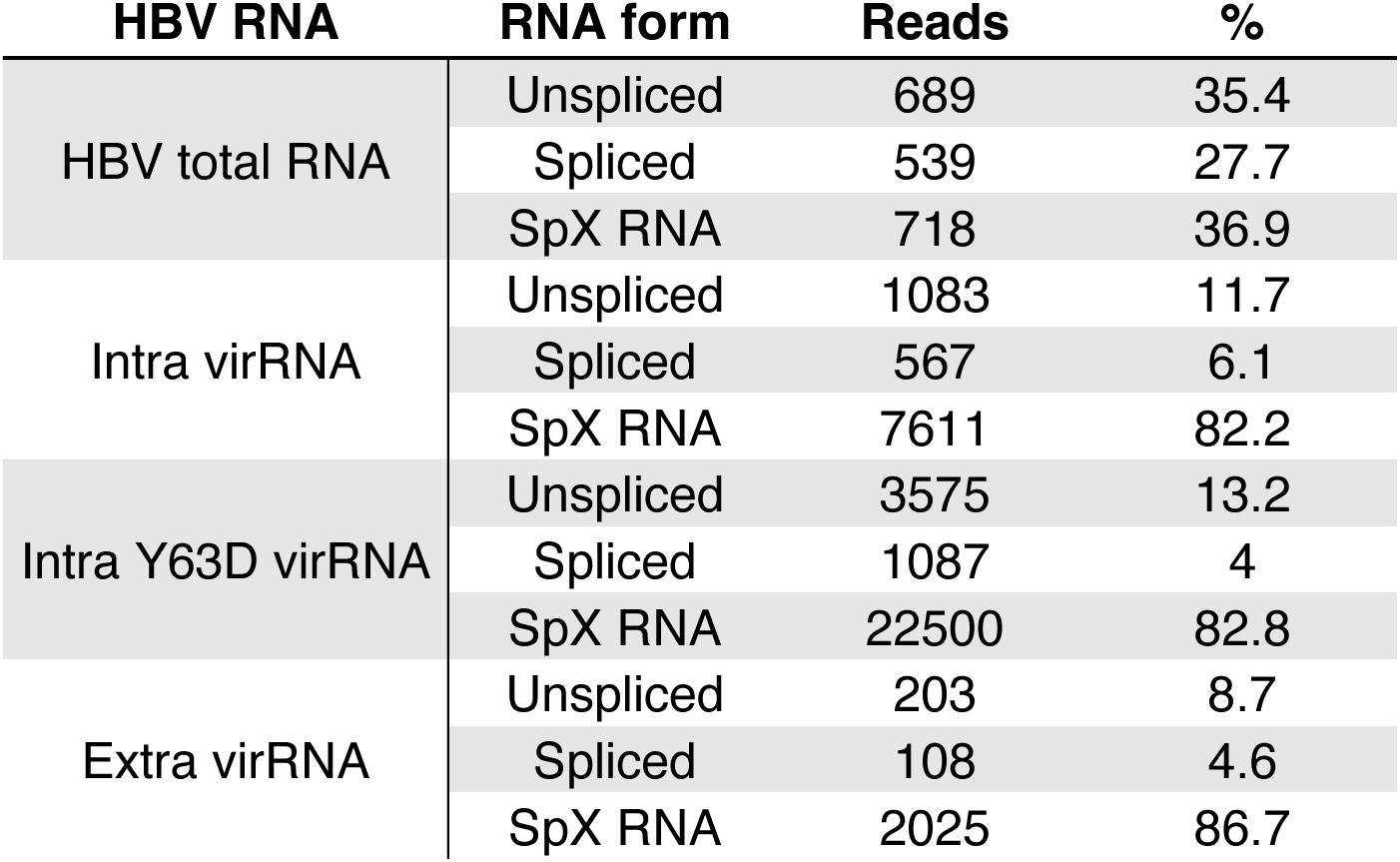
Proportion of HBV SpX isoform by Nanopore DRS.

